# The Bayesian Superorganism: externalised memories facilitate distributed sampling

**DOI:** 10.1101/504241

**Authors:** Edmund R. Hunt, Nigel R. Franks, Roland J. Baddeley

## Abstract

A key challenge for any animal (or sampling technique) is to avoid wasting time by searching for resources (information) in places already found to be unprofitable. In biology, this challenge is particularly strong when the organism is a central place forager – returning to a nest between foraging bouts – because it is destined repeatedly to cover much the same ground. This problem will be particularly acute if many individuals forage from the same central place, as in social insects such as the ants. Foraging (sampling) performance may be greatly enhanced by coordinating movement trajectories such that each ant (‘walker’) visits separate parts of the surrounding (unknown) space. We find experimental evidence for an externalised spatial memory in *Temnothorax albipennis* ants: chemical markers (either pheromones or cues such as cuticular hydrocarbon footprints) that are used by nestmates to mark explored space. We show these markers could be used by the ants to scout the space surrounding their nest more efficiently through indirect coordination. We also develop a simple model of this marking behaviour that can be applied in the context of Markov chain Monte Carlo methods (Baddeley et al. 2019). This substantially enhances the performance of standard methods like the Metropolis–Hastings algorithm in sampling from sparse probability distributions (such as those confronted by the ants) with little additional computational cost. Our Bayesian framework for superorganismal behaviour motivates the evolution of exploratory mechanisms such as trail marking in terms of enhanced collective information processing.

## Introduction

Exploring an unfamiliar, changing environment in search of valuable resources such as food or potential nest sites is a challenge for many organisms. A memory of where one has already explored, to avoid revisiting unprofitable locations, would generally seem to be an advantage. A spatial memory of foraging locations, for example, is likely to be beneficial but it would entail physiological costs. These include the metabolic overhead of a bigger brain (memory storage capacity) and the cost of encoding and retrieving memories (brain activity); these costs have to be traded-off against the benefit of improved foraging performance [1]. One way to circumvent the cost of carrying memories internally is to store the information externally in the environment. Indeed, such externalised information storage may have been the historical precursor to the development of internalised memory storage and retrieval [2,3]. Use of external markers as memory has been demonstrated in arguably simple organisms, including the slime mould *Physarum polycephalum* [4]. Markers may be left in the environment to signify the presence or absence of good foraging or nest-making prospects at that location, so that when an animal returns it can make an appropriate and timely decision about the expenditure of its efforts.

An analogous problem to animals searching for food sources in an unfamiliar environment is that of sampling efficiently from unknown probability distributions. Markov chain Monte Carlo (MCMC) methods have been developed to generate such samples but suffer from the same problem of potentially revisiting the same area of probability space repeatedly. Although methods such as the Metropolis– Hastings algorithm show random walk behaviour in their sampling trajectory, they are still popular techniques because of their ease of implementation and computational simplicity. As discussed in the first paper of our series on the ‘Bayesian superorganism’ [5], MCMC methods can be used as a model of animal movement that enacts the optimal probability matching strategy for collective foraging (‘gambling’) [6]. There, we drew a parallel with advances in efficiency in MCMC methods, to more adaptive animal behaviours emerging in natural history: for example, from the random-walk type behaviour of the Metropolis-Hastings algorithm [7,8] to the correlated random walks produced by Hamiltonian Monte Carlo [9]. The performance boost provided by externalised memories, discussed here, may similarly be understood as an evolutionary advancement in collective information processing capabilities.

Here, we present two main findings: (1) evidence that *Temnothorax albipennis* worker ants avoid the footprints of nestmates during the reconnaissance of an unfamiliar space, and that this enhances the efficiency of the colony-level collective exploration; (2) we develop a bio-inspired trail avoidance method for the purpose of Markov chain Monte Carlo sampling that similarly uses a memory of where the walker has visited to obtain a representative sample more rapidly. This method is easy to implement and computationally simple, and offers significant improvements in performance for the sort of sparse distributions that ants and other organisms also have to sample from routinely. In the next section we introduce some more biological background on chemical markers (externalised memories), but the reader familiar with this topic may proceed to the Methods section.

### Further biological background on chemical markers

In addition to expediting the process of environmental search, external markers can also communicate one’s presence to friendly or competitor conspecifics. Some animals are well-known for marking territory: for instance wolves (*Canis lupus*) use scent marks to maintain their territories [10] while mice (*Mus musculus*) use olfactory cues to help establish their territorial boundaries [11]. In the Hymenoptera, the honeybee *Apis mellifera ligustica* marks visited flowers with a scent and rejects flowers that have been recently visited; it responds more strongly to its own markers than that of its nestmates [12]. Such repellent scent-marks should help the bee to forage more efficiently. As in the ants, these marks may simply be hydrocarbon footprints rather than costly signals [13]. Ants are known to engage in two distinct approaches to marking the area around their nest. In territorial marking, ants will defend the marked area against intra- and inter-specific intruders. By contrast, home-range markers label areas that the colony knows to be hospitable and available for foraging, but such areas are not defended against other ant colonies [14]. Major workers of the African weaver ant *Oecophylla longinoda* mark their territories with persistent pheromones that are distinguishable to the ants at the colony level [15], whereas workers of the leafcutter ant *Atta cephalotes* deposit ‘nest exit pheromones’ in the vicinity of their nest entrances, helping to orient workers and hence enhancing the efficiency of leaf gathering [16]. *Lasius niger* ants have been found to engage in home-range marking, which rather than territorial marking may help them to enlarge their potential foraging space alongside other colonies, in a ‘shared information’ strategy [17].

Apart from pheromones, which are signals laid deliberately by ants at a cost to themselves, to influence the behaviour of other ants, ants have also been shown to respond to cues such as the cuticular hydrocarbon footprints left by other ants. These can also be used to distinguish between different colonies [18], and potentially also to recognize different castes, as in a study of *Reticulitermes* termites [19]. They can also be detected by ant predators, and so this deposition is not without costs [20]. The response ants make to detecting such markers depends on whether the cue is from a nestmate, a competing colony, or a different species altogether, because depending on the relationship between sender and receiver, it could represent a threat or useful information about the location of foraging resources [21]. The role of the ant species (dominant or submissive) in the local community can be key in this respect [22].

In our study organism, the house-hunting rock ant *T. albipennis*, there is evidence for pheromone use in navigating to a new nest site [23,24] as well as in *T. curvispinosus* [25]. Other *Temnothorax* species are also known to use individual-specific trail pheromones for orientation outside their nests: *T. affinis* [26] and *T. unifasciatus* [27]. *T. albipennis* is thought to use an individual-specific pheromone trail during the nest-site assessment process: to measure the floor area using the ‘Buffon’s needle’ algorithm, an ant experiences a lower crossing rate of its own trail when the nest is larger. If its trail was not individual-specific only one ant could do this before the trail was confused with those of other scouting nestmates [28]. *T. albipennis* colonies are also found to avoid conspecific colonies when choosing where to locate its nest, as a result of odour cues left by foreign colonies [29]. Pheromones are also probably used by the ants when scouting potential new nest sites, since when previously assessed nest sites are replaced with clean nest sites, colonies are more likely to pick the poor option, having lost their reconnaissance information [30]. *T. rugatulus* was found to use an alarm pheromone that elicits two different behaviours depending on the context: in an unfamiliar nest it causes ants to reject it as a potential new home, to avoid danger; while when released near the ants’ home nest it attracts them to defend against a threat [31].

Chemical marks left by harvester ants (*Pogonomyrmex*) during exploration for new food have been suggested as a way of avoiding redundant repeat searches in that area [32,33]. The usefulness of an avoidance behaviour during the searching phase for food was commented upon by Nonacs in a study of *Lasius pallitarsis* ants. Such a behaviour was predicted to be more helpful for solitary foragers, than for species with mass recruitment to prey items that need teams of workers to carry them home. Experimental investigation found that the ants show a state-dependent avoidance of the path of previous nestmates [34]. This question has recently been revisited when evidence was found that *Myrmica rubra* (Myrmicinae) ants avoid the footprints of their nestmates, which suggests that they are using the information it conveys to inform their foraging or scouting decisions [22]. However, this behaviour was not observed in the four other ant species in the study: *Formica polyctena* (Formicinae), *Formica rufibarbis* (Formicinae), *Lasius niger* (Formicinae), and *Tetramorium caespitum* (Myrmicinae). *T. albipennis* is also in the subfamily Myrmicinae, and we predicted that it too uses cuticular hydrocarbon footprints left by previous exploring nestmates to scout an unknown area surrounding its nest more efficiently. Here, we present evidence that *T. albipennis* indeed avoids the footprints of nestmates during the reconnaissance of an unfamiliar space, and for the first time show that it enhances the efficiency of the colony-level collective exploration.

## Methods

### Data processing and analysis

We examine data from a previous study of *T. albipennis* ant movement patterns [35,36]. These data present two experimental regimes, one in which the foraging arena was entirely novel to exploring ants, because it was cleaned by the experimenter in advance, and one in which previous traces of the ants’ activities remained. We call these treatments ‘No cleaning’ (NC) and ‘Cleaning’ (C). For both treatments, we used 18 ants, 6 from each of the three colonies, for a total of 36 ants. Treatments were alternated over six consecutive days. The colonies inhabited small artificial nests with a 2mm diameter entrance hole drilled into the centre of the top. A colony was placed into a 90×90cm arena and a 15×15cm white paper mask with a 1×1cm central hole affixed over the top, with a further 1.5×1.5cm white paper removable cover to block the nest entrance. This created a smooth surface (with a raised area over the nest) to allow continuous tracking of a single ant. The paper cover was removed to allow a single ant out of the nest, and the entrance cover was then replaced, so that the ant would explore freely in isolation for 45 minutes before being removed to a separate dish for the duration of the experiment. The exploration trajectory of the ant was recorded by a camera mounted on a motorised gantry system, which followed the ant’s movements and recorded its path as a sequence of (*x*, *y*) points spaced by 0.1 seconds. In the NC treatment, six ants were consecutively released from the same colony on the same day, and pheromone signals and/or other cues such as cuticular hydrocarbon footprints were allowed to accumulate. In the C treatment, the arena surface was cleaned before the subsequent ant was allowed into the arena, to prevent chemical communication between successive ants. See the ‘Methodology and apparatus’ section of Hunt et al. [36] for a more detailed description of methods and Basari et al. [37] for a photo and more details on the gantry system.

In the previous study [36] the focus of the analysis was on the characteristics of movement ‘events’ – a period of non-zero speed followed by stopping – and the effect of treatments on metrics such as average speed and the correlation between the average speed of consecutive movement events. Here, we examine the effect of the treatment on exploration behaviour – how efficiently the ants reconnoitre an unfamiliar experimental arena – with the hypothesis that the ants use chemical information from previous exploring nestmates to do this more quickly, by avoiding moving through regions of the space that have already been somehow marked as explored. We test this hypothesis of enhanced exploration efficiency in the following way. In nature the ants might benefit from sampling preferentially from surrounding regions of higher quality, in the sense of being more likely to contain valuable resources such as food [5] or a potential new nest site [38]; and would want to spend less time in ‘empty’ regions of the space that contain little of relevance to their reproductive fitness. High-quality regions are likely to be associated with cues that help the ants navigate toward them: for instance, a good potential nest site may be more likely to be located in brighter regions of space than darker regions, because it could benefit from warmer ambient temperatures. Therefore, in a natural environment we would not expect the ants to explore uniformly in the region around their nest; and while we might naively suppose that an empty experimental arena represents a uniform environment, from an ant’s perspective it may be somewhat heterogeneous (lighting, air currents, vibrations, magnetic field, temperature, thigmotaxis around the edges of the paper mask with a focus on the corners, and so forth), despite an experimenter’s best efforts. Therefore, rather than measure the ants’ exploration efficiency by reference to how quickly they fill uniform space, we take the ‘target distribution’ that ants are trying to sample from as being approximated by the actual exploratory trajectories taken by all 36 ants. While this combines data from two different treatments, with social information presumably present in one (‘no cleaning’), this larger sample is more likely to approximate the ants’ exploration preferences. Although this ‘target distribution’ may seem an unconventional approach, it will account for any heterogeneity in terms of the lighting environment, for example (though best efforts were taken to control conditions), and also differential responses in terms of lateralized behaviour. For instance, a preference for viewing landmarks with the right eye has been suggested [37], which may affect exploration behaviour, and also a leftward turning bias when exploring nest sites [39], which may relate to these ants having slightly asymmetric eyes [40].

We create a target distribution for the exploring ants in the following way, using MATLAB 2015b. The exploration trajectories for all 36 ants are combined (a total exploration time of 27 hours), and trajectory points located inside the central region where the 15 × 15 cm paper mask is located are removed, because many of the ants spend a significant proportion of their time familiarising themselves with their immediate surroundings and engaging in thigmotaxis (edge following) around the mask before beginning reconnaissance bouts away from the central region. The (*x*, *y*) coordinates were then transformed from a Cartesian to a log-polar system (*θ*, ln *r*), where −*π* ≤ *θ* ≤ *π* and 4.3 < ln *r* < 6.5 (the approximate size of the arena in mm excluding the central area). This transformation of trajectories makes the distribution of visits to different parts of the arena more uniform, because a spatial histogram effectively has larger bins for more radially distant areas, and smaller bins for the more densely visited central region containing the nest. This makes statistical tests for differences between trajectories more powerful. A periodic boundary for angle *θ* was created by copying the points in the rectangular region *θ* ∈ [−*π*, 0], ln *r* ∈ [4.3,6.5] into a new region [*π*, 2*π*] and similarly copying [0, *π*] into [−2*π*, −*π*] to create a larger space −2*π* ≤ *θ* ≤ 2*π*. This periodic boundary prevented ‘ringing’ effects at the edge when a gaussian blur was applied over all the points in the space, to create a continuous distribution. This also functions effectively as data interpolation for regions without data, because the small ants (1mm wide) will not visit every point in the 900 mm square arena. The coordinates from all 36 trajectories were counted using a 2D histogram on a grid 1257 × 881 (*θ* step size 0.01, ln *r* step size 0.0025). This represents where in the arena the ants tend to spend their time exploring. A gaussian blur has a parameter *σ* which controls the width of the blur. Since the ants are small (only around 1mm wide) a fairly wide blur is necessary to transform their exploration trajectories into a continuous distribution. The *σ* used was set equal to 30, which corresponds to larger blurs for more radially distant (less visited) regions when applied in ln *r* space: roughly 6mm wide closest to the nest and 40mm wide toward the edge of the arena. After the blur was applied to the 2D histogram, the distribution was then truncated back to its original size such that −*π* ≤ *θ* ≤ *π* again, and normalised such that the sum of all the points was equal to 1, as in a discrete probability distribution where the value at each arena grid position represents the probability of an ant visiting it. The ants’ exploration trajectories and resultant empirical target distribution is shown in Figure 1, and individual and cumulative trajectories are shown in Figures S3-6.

**Figure 1.**
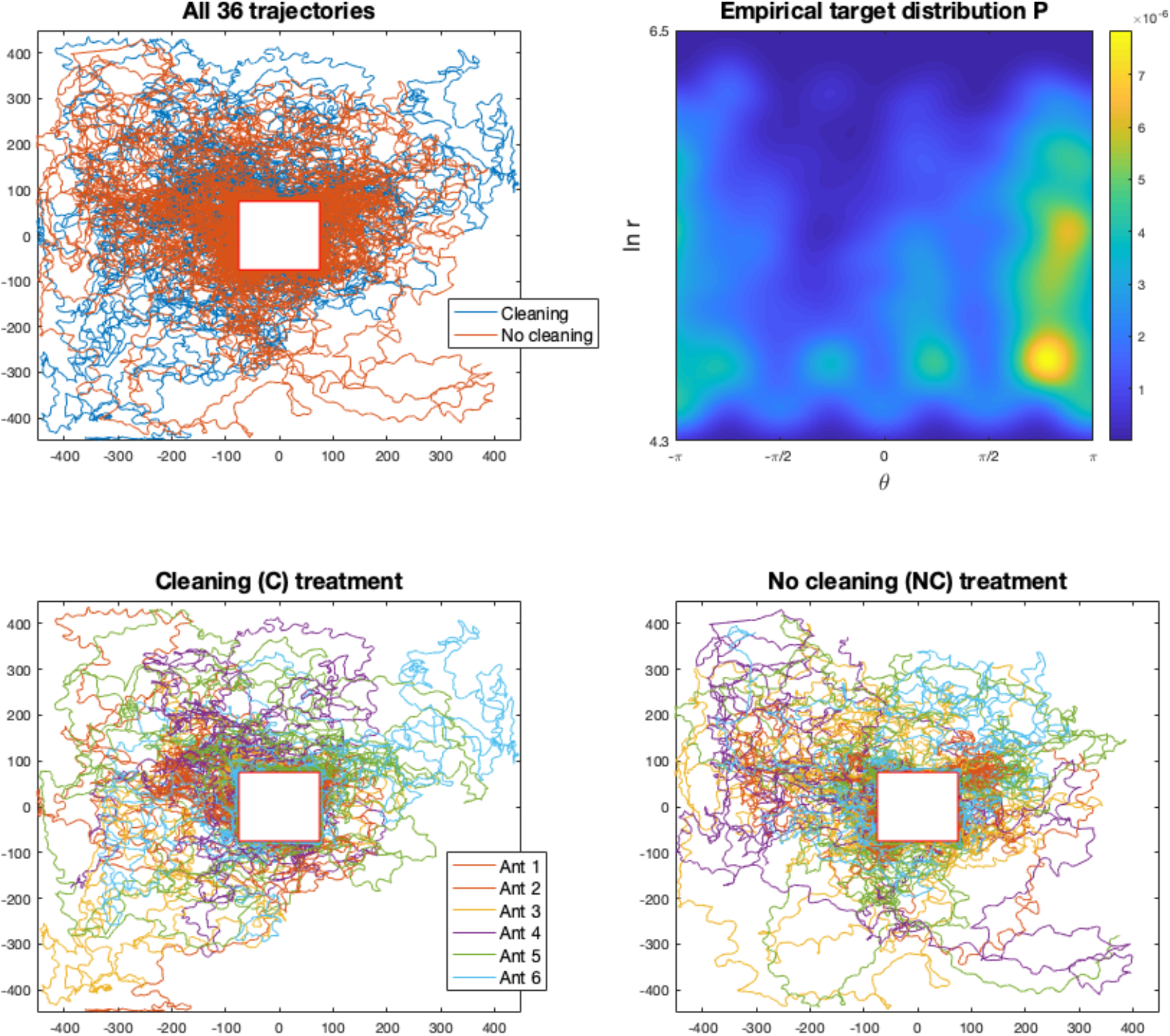
The ants’ exploration trajectories for the two treatments and empirical target distribution P. All trajectories are shown in the top left pane, marked by treatment, while trajectories are shown by treatment and ant order in the bottom panes (3 ants for each order in the sequence).

As shown in Figure 1, the ants in the cleaning treatment made a somewhat uneven exploration of the arena, spending more time in the top-left (quadrant 2) and not visiting the bottom-right corner of the space (quadrant 4), in contrast to the ants in the no cleaning treatment. We sought to compare our data with an independent study of a different ant species to investigate whether such exploration heterogeneity had precedent elsewhere. We examined the data of Khuong et al. [41], which recorded the exploration behaviour of *Lasius niger* ants placed individually in an empty arena of various inclines. Their treatment with zero inclination recorded the movement of 69 ants from the centre of a 50 × 50 cm arena to the edge, whereupon the ant was removed. It should be noted that these worker ants are larger at around 3-5mm long compared to 2mm *Temnothorax* ants, and so our arena was larger from the ants’ perspective as well as in absolute terms. We examine which side of the arena the *L. niger* ants exited the 50×50cm region and replicate this analysis for our own ants (Figure S1). Although the exploration of the ants was uniform in the centre of the arena, it had a bimodal angular distribution as the radius of exploration increased (Figure S2). We perform a Chi-Square goodness of fit test for a uniform distribution of exit sides (i.e. an expectation of 25% on each side), and an exact multinomial test using the R package ‘EMT’ [42] for our smaller number of observed exit points. Because the first ants in the NC treatment experience a clean arena, we switch these three ants to the C grouping for the purpose of this analysis.

To measure how quickly each treatment converged toward the final distribution, the exploration trajectories were combined by colony from all ants up to ant 1,2, … 6 out of the nest. For instance, to measure the progress by ant 2 of the NC treatment, the trajectories of 2 ants were combined (the first ant, and the second) to see its cumulative progress toward the goal. The process described above was then repeated to obtain a probability distribution, which was then compared against the final distribution using the cross-entropy (CE) as a measure of their similarity, where a lower CE signifies greater progress toward the equilibrium distribution of a well-explored arena (see also [5]). In the last step the three colony-level CE measures were averaged to obtain a mean cross-entropy for the NC treatment by ant 2. Our hypothesis that the ants use chemical information from preceding nestmates to explore more efficiently would correspond to a CE that falls faster in the NC treatment than the C treatment.

While the NC and C treatments measure the difference to exploration made by chemical information from nestmates, it is also quite possible that individual ants make use of their own pheromone markers or other cues left behind, like their own footprints, to avoid wasting time revisiting regions of space they have already walked through. To assess the benefit of individual-level memory (both internal and external), we constructed simulated trajectories by randomly sampling from a vector of the movements of all 36 ants. These ‘Markov’ ants provided a memoryless benchmark of exploration performance, as ants that do not avoid their own trails, because there is neither externalised memory (chemical markers) nor internal memory (in their brain) of where they have been. A trajectory of 45 minutes of exploration was constructed by concatenating 1 second periods of movement that stayed within the arena boundary (a 90×90cm space). The performance of 100 ‘colonies’ of these Markov ants (6 per colony, as with the real colonies) was compared in the same way.

To assess the statistical significance of one treatment converging more quickly on the final distribution than the other, a permutation test was used, on ants 2-6 out of the nest. The first ant is not included in the test for statistical significance since it is expected that the two treatments are equivalent at this stage: the first ant explores an arena empty of any chemical information. A non-parametric test is appropriate since the distribution of the CE for each treatment is unknown, and relatively few data points are being compared (15 from each treatment). A permutation test was also used to assess whether the average distance travelled by each exploring ant is different in the NC and C treatments.

### Trail avoidance model

In previous research [5,38] we have proposed that a central challenge that an ant colony confronts, that of trying to identify regions of value in an unknown surrounding environment, is analogous to the problem found in statistics, physics, etc. of efficiently sampling from complex probability distributions or computing numerical integrations. This is because in order to sample from a probability distribution efficiently, one must identify regions of high probability density, but these are not known *a priori* but only after evaluating the probability density function at those locations. Similarly, an ant needs to find regions of high quality, but does not know where they are unless it expends effort (time, energy) in exploration [5,38]. An ant colony has a key advantage over a solitary animal: it is composed of many workers, which can cooperate to benefit the colony as a whole, working together as one ‘superorganism’ [43]. Each worker could set off in a different direction to that of its nestmates, and hence all of the surrounding area could be quickly explored. However, the ants lack any central controller to direct their movements so as not to overlap and waste time exploring the same region several times; instead they must make their decisions about where to move based only on locally available information [44]. A key feature of the Markov chain Monte Carlo (MCMC) movement models we have proposed is the use of local quality gradient information in deciding where to move next: it would seem reasonable to expect ants (and other organisms) preferentially to move toward regions of increasing quality [5]. What we have not yet considered is that this gradient may be modified by the ants themselves, by making use of their sophisticated chemical communication systems [14]. We suggest that ants may leave behind chemical markers, either deliberately (i.e. pheromones) or incidentally (such as hydrocarbons from their footprints), that convey information about where they and their nestmates have already explored. Such markers may be thought of as ‘negative’, in the sense that an ant seeking to explore the environment will seek to avoid them, and as reducing the quality gradient in that direction, such that other alternative directions have a relatively higher gradient.

In a simple encapsulation of this exploration and trail avoidance behaviour, we can say that an ant trying to sample from an unknown resource quality distribution *P* also leaves behind a consistent marker at every step that it takes, which corresponds to its subjective experience or model of the world *M* (but which does not require any internal memory in the ant itself). *M* starts off as a uniform prior of the world (e.g. a distribution of ones for each location) but then is added to (e.g. +1) at each visited location. This is equivalent to Bayesian updating with a Dirichlet prior, because this is the conjugate to the multinomial distribution [45]. *M* can be renormalised at each time step to be a probability distribution that sums to 1. Over time, as the ant more thoroughly explores the distribution *P*, this model will converge *M* → *P*, with more markers accumulating in regions of higher probability. Since the objective of the ant is to sample from regions of high probability first, one way to prioritise such regions is to sample from *P*^2^, since this will substantially increase the relative value of high probability regions over low probability regions. For instance, an ant would rather visit a region of 0.2 probability over a region of 0.1, with the former being of 2 times the quality as the latter. If this is squared, this becomes a relative value of 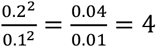 times higher, making this region of P space much more attractive, with a sharper gradient toward it too. Therefore, an effective objective for ant exploration (or indeed MCMC methods) is to sample from:

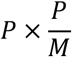

which converges in time to sample from *P* as *P*/*M* → 1, but which should sample from high probability regions first as a combination of the *P*^2^ (attractive) and *M* (repulsive) terms. Such an objective function for the ant or MCMC method to sample from does not require a change in the method itself: it can be applied in conjunction with the most basic approaches such as Metropolis–Hastings, which are computationally simple. A ‘pseudocode’ representation of our approach is shown in Table 1.

**Table 1.**
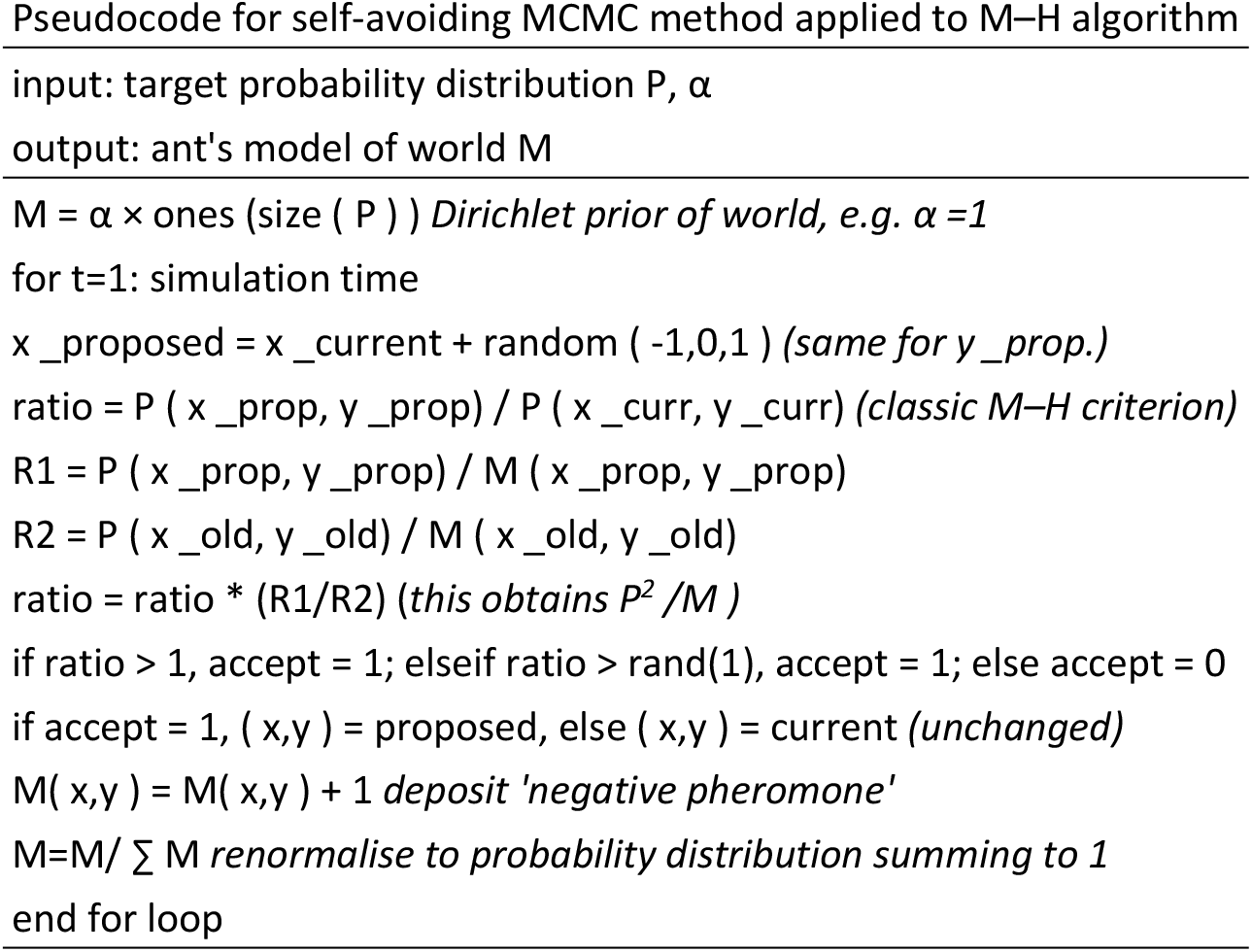
Pseudocode for the self-avoiding MCMC method.

Simulations of this ‘trail-MCMC’ model of ant exploration (or sampling approach) are presented in the results, using a Metropolis–Hastings (M–H) method to sample from both a random distribution (corresponding to a noisy, sparse world) and the empirical distribution of all the ant trajectories. The random distribution (shown in Figure 4) was Dirichlet-distributed, generated by sampling from a gamma distribution of shape *k* and scale parameter *θ* = 1. This results in mostly low-probability cells with a few higher-probability cells, which corresponds to the ants’ ecology where most space is unsuitable for locating the colony’s nest.

The M–H method samples from the distributions using periodic boundary conditions. To assess the effect of cleaning (that is, removing the externalised spatial memory of the ants) the simulations are also run under a ‘cleaning’ condition where the memory is reset five times, to correspond to the five cleaning bouts in the experimental study (after each ant has explored, for six ants). To ease this comparison the simulations are run for T=60,000 time steps, or 10,000 time steps per ‘ant’. The performance is compared to a M–H method sampling from the standard objective distribution *P* by examining the rate of fall in cross-entropy between the sample and the target *P* (as previously discussed in [5]).

## Results

### Ant exploration trajectories

The rate of convergence of the exploring ants toward the final distribution is shown in Figure 2; the ants in the no cleaning treatment converge significantly more quickly than those in the cleaning treatment (p=0.0161, permutation test on ants 2-6 across the 3 colonies. The analysis was repeated for Gaussian blur size *σ* = 20, resulting in p=0.0394, and *σ* = 40, resulting in p=0.0098). This is despite the average distance travelled by the ants from each treatment not being significantly different. The average distance is 18.58 ± 0.77 m (s.e. of the mean) in the NC treatment and 17.57 ± 0.77 m in the C treatment (p=0.21, permutation test). As expected, without any memory (externalised or internal) the simulated Markov ants converge on the target more slowly than either of the experimental treatments, because they are essentially engaged in a random walk around the arena.

**Figure 2.**
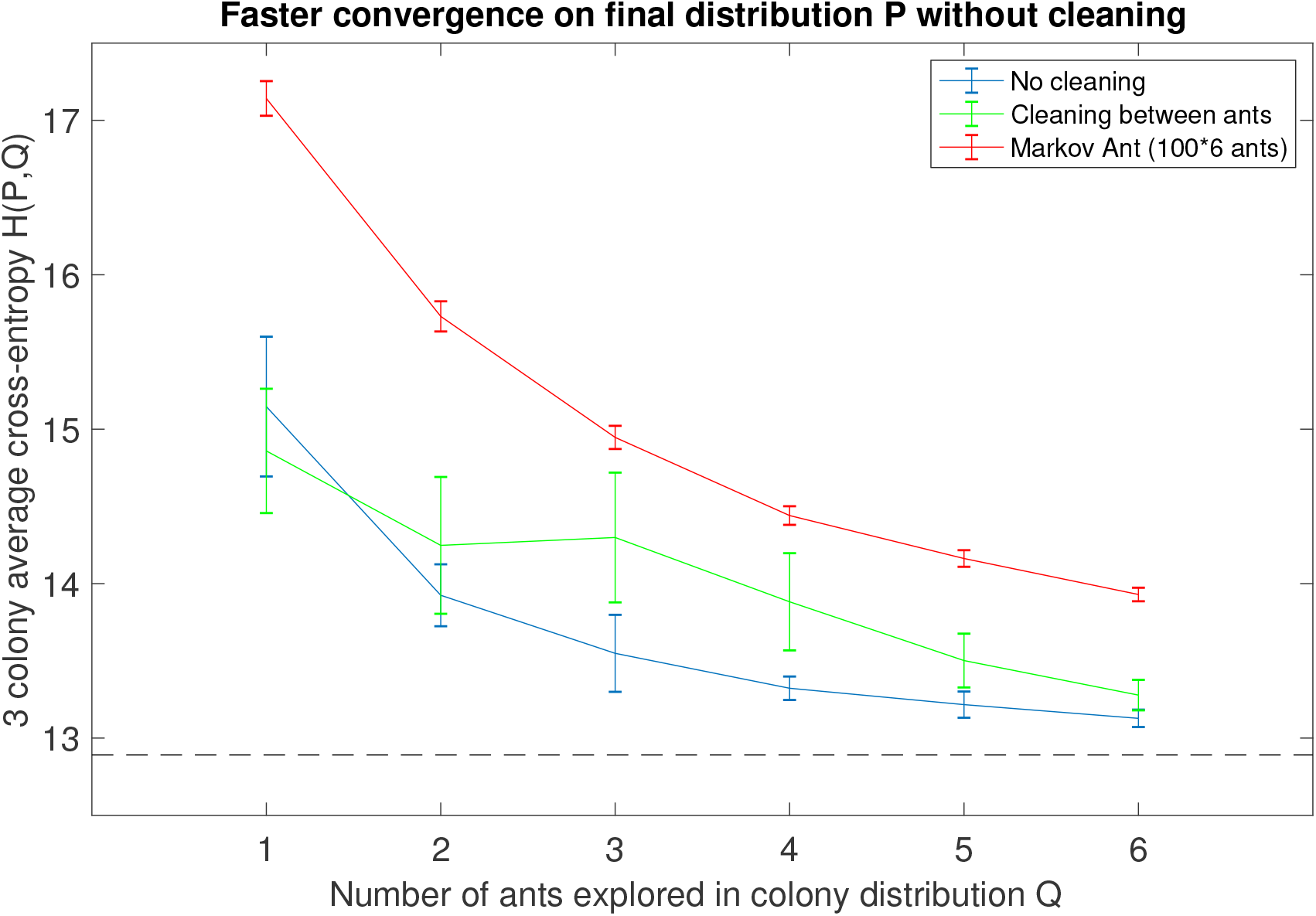
The ants in the no cleaning treatment converge most quickly toward the final distribution, indicating that they benefit from chemical information left by preceding nest mates. The error bars indicate one standard error of the mean; a permutation test on ants 2-6 indicate that the cross-entropy is significantly lower in NC than C (p=0.0161). The simulated ‘Markov ant’ converges slowest, because it benefits neither from the chemical markers (externalised spatial memory) nor internal memory about where it has already visited. The cross-entropy minimum is indicated by the dashed black line.

As anticipated, the ants were not uniform in how they explored the arena. Ants in both treatments tended to spend more time in the second quadrant of the arena and less time in quadrant 4 (Figure 1); this could represent a tendency in the ants to respond to unknown environmental gradients, or some kind of intrinsic behavioural tendency. Ants in the no cleaning treatment tended to cover more of the arena as a whole (Figure 1). We confirmed that heterogeneity in exploration patterns is not unique to our data by examining the *Lasius niger* data of Khuong et al. [41] which showed a significant tendency for ants to leave through two of the four virtual arena sides (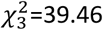, p=1.384×10^−8^, Figure S1). While there was a significant exit bias for ants in our cleaning treatment (exact multinomial test: p=0.0137), this tendency was not found to be significant in the no cleaning treatment (EMT: p=0.1102).

### Trail avoidance model simulations

An example simulation of the models (standard M–H, trail-avoidance M–H with and without cleaning) for the sparse distribution is shown in Figure 3. Such a distribution corresponds to a region of empty space where the ants would wish to avoid sampling from the same parts of the distribution repeatedly. It might be thought of as a ‘zoomed-in’ patch of empty space in the ants’ environment, as opposed to the more patchy empirical distribution in Figure 4. Figure 3 shows that the standard M–H method converges slowly to the target distribution when all the space needs to be sampled from roughly equally: with a random walk considerable time is wasted going over previous ground. By contrast, the trail-avoidance M–H methods converge much more quickly, as seen in a more rapidly falling cross-entropy. In a way similar to the experimental result shown in Figure 2, the advantage of not cleaning the pheromone trail is not immediately apparent but becomes evident by ‘ant 3’ (*t* = 3 × 10^4^ on the *x* - axis), because more pheromone has been accumulated for the no cleaning ant to avoid. In both Figure 3 and Figure 4, all methods would eventually converge to the target distribution given enough time: but enhanced sampling efficiency has significant value for both biological fitness and technological applications.

**Figure 3.**
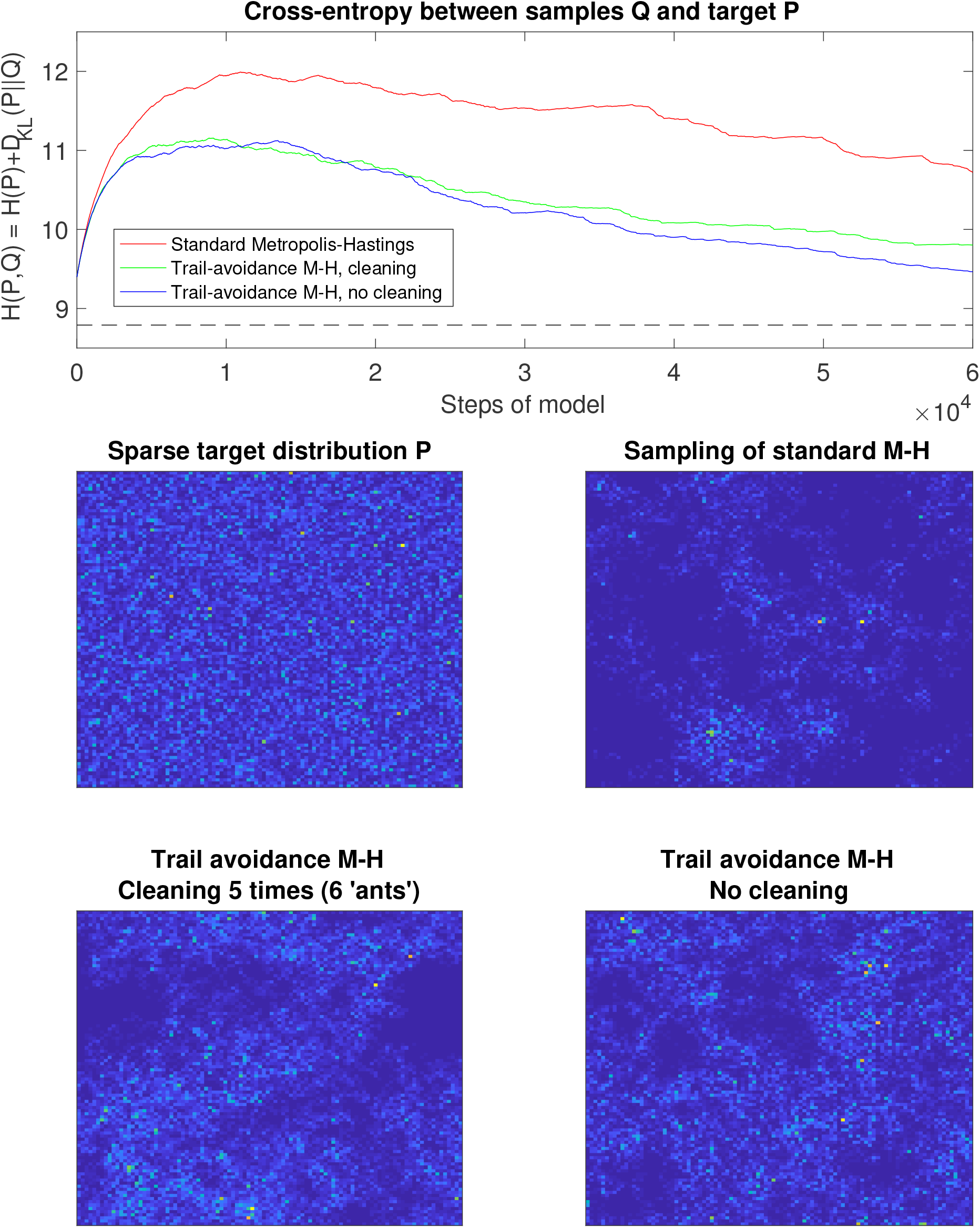
The trail avoidance model converges much more quickly to the simulated sparse target distribution, while ‘cleaning’ (removing the externalised memory of the sampling trajectory) slows its progress somewhat. The target (cross-entropy minimum) is shown by the dashed black line.

**Figure 4.**
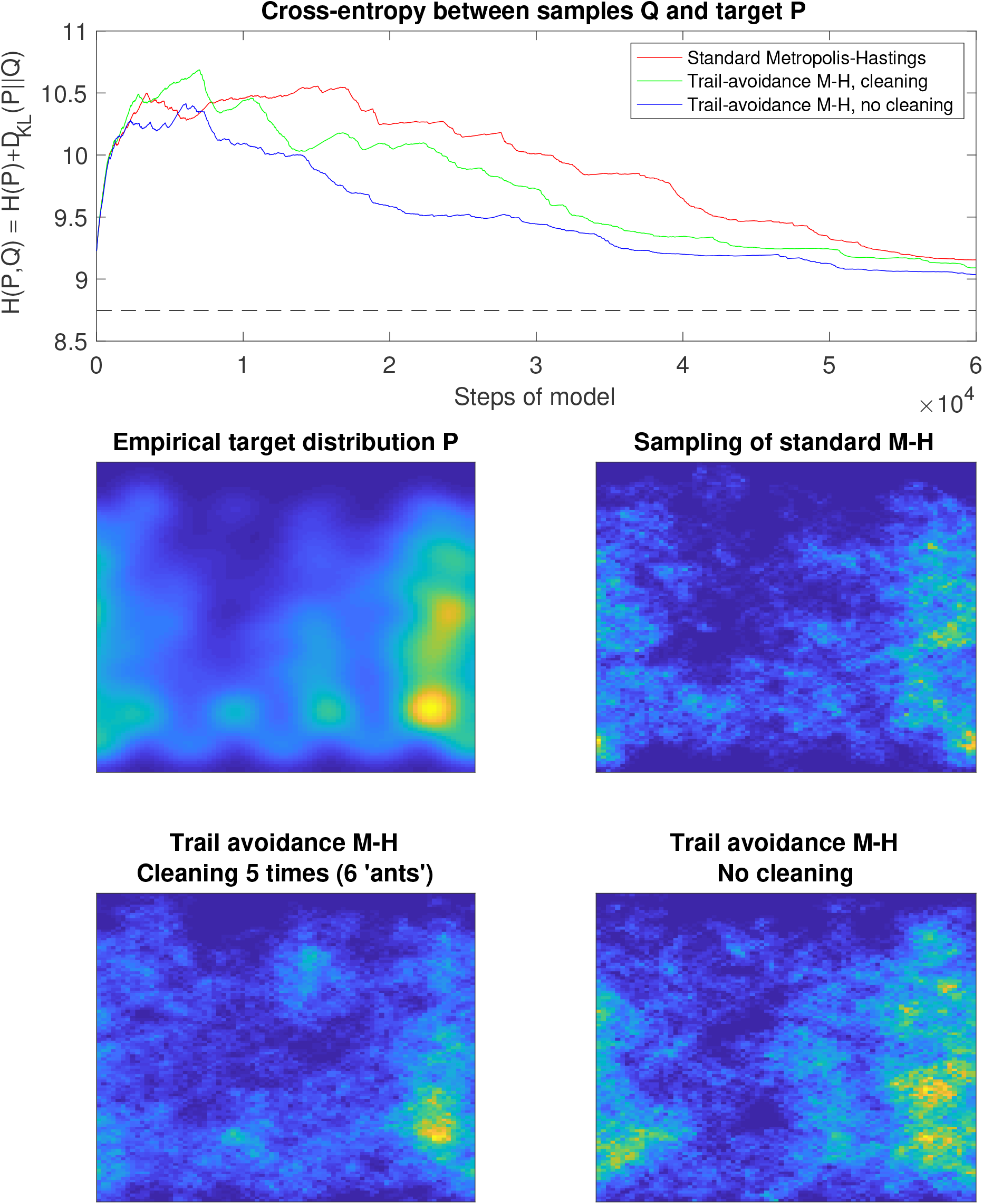
The trail avoidance model again converges more quickly to the target. The M–H model is more likely to get ‘stuck’ in higher probability regions and take longer to get to other important parts of the distribution. However, the performance gain is less obvious than in the preceding gamma distribution example.

The result of a set of simulations sampling from the empirical distribution of Figure 1 is shown in Figure 4. This does not show such a large improvement in performance for trail-avoidance M–H over standard M– H, because significant portions of the probability mass are more readily accessible to random walk-like behaviour. Nevertheless, the standard M–H method is seen to have spent more time in the quadrant 2 patch, in keeping with the general problem the ants also face of prematurely favouring a local resource, rather than exploring relevant information in the global distribution. In comparison, the no cleaning trail-avoidance M–H samples from the whole distribution in proportion to their quality in this example, and converges most quickly to the target distribution as a result. The cleaning simulation shows intermediate performance, sampling more effectively than M-H but not exploring the whole distribution as thoroughly as when the pheromone memory is allowed to accumulate.

## Discussion

Ants and other organisms confront the challenge of having to explore unknown environments while avoiding wasting time and incurring other costs and risks, which would occur if there are repeated visits to the same unprofitable areas. One strategy to avoid time-wasting would be for the animal to maintain an internal memory of where it has been, by using techniques such as learning the configuration of local landmarks (allothetic navigation) [46], or counting steps in a certain direction from the nest (path integration, or other idiothetic mechanisms such as optic flow). However, the ants are pre-eminent manipulators of terrestrial chemical information [14]. Analysis of experimental data [35] presented here provides initial evidence that *Temnothorax albipennis* ants use chemical depositions – either pheromones or cues such as cuticular hydrocarbon footprints – to explore an unknown arena more efficiently. This is because the ants in the treatment where chemical information from nestmates is allowed to accumulate converge on the colony’s target distribution more quickly than when that information is systematically cleaned away. Such indirect coordination between ants is consistent with previous analysis of this data, which found that the correlation between the speeds of successive movements was significantly lower in the ‘no cleaning’ treatment (0.234 vs. a correlation of 0.407 in the ‘cleaning’ treatment), which we suggested is because ants are responding to social information by changing their movement patterns [36]. Such colony-level indications of a potential negative marker could be confirmed in future research by using coupled gas chromatography/mass spectrometry to identify candidate marker compounds from arena swabs in the ‘no cleaning’ treatment. The influence of such candidate compounds could then be tested experimentally on individuals or colonies, as in Sasaki et al., for example [31].

Our finding of collective exploration complements previous research on another Myrmicine ant, *Myrmica rubra*, that found evidence for nestmate footprint avoidance though did not demonstrate enhanced search efficiency [22]. The avoidance behaviour is seen to be such that the trajectories of two successive ants will nevertheless intersect several times: persistent avoidance would, in effect, paint an ant into a corner. Instead the suggestion of a previous ant’s reconnaissance leads to a reweighting of exploration toward a different region of space. In practice, where an ant holds private information about a source of food, for instance, that conflicts with social information (an exploratory marker) it is likely to dominate the decision about whether to return to the space when it is held with high confidence (probability) [56]. When there is a coherence between private memory and social information, this may help to boost ants’ ‘confidence’ in their own memories and walk more directly (straighter and faster) toward their goal [57].

The results shown in Figure 2 indicate the value of both internal and external forms of memory: the ‘Markov ants’ have neither, while the ants in the two treatments (cleaning and no cleaning) have both available, but the no cleaning ants to the fullest extent. It may be possible to assess the relative contribution to exploration efficiency from own versus others’ pheromones by covering an exploring ant’s gaster with paint [54], while the use of visual landmarks to navigate [46] can similarly be prevented by covering an ant’s eyes [55]. It may be that ants benefit primarily from their own chemical marks and only secondarily from nestmate marks. These experimental possibilities are summarised in Table 2, and could be used to quantify the value of different information sources for exploration, in informational terms, using the cross-entropy method introduced here.

**Table 2.** Additional conditions that could be investigated to assess the value for exploration of different information sources.

| Condition | Internal Memory |  | External Memory |  |
| --- | --- | --- | --- | --- |
|  | Other<br>(odometry etc.) | Visual | Pheromones |  |
|  |  |  | Own | Others |
| Markov ant | x | x | x | x |
| Covered eyes | ✓ | x | ✓ | ✓ |
| Covered gaster + cleaning | ✓ | ✓ | x | x |
| Covered gaster | ✓ | ✓ | x | ✓ |
| Cleaning | ✓ | ✓ | ✓ | x |
| No cleaning | ✓ | ✓ | ✓ | ✓ |

There was a noticeable tendency for ants to explore quadrant 2 of the arena in preference to quadrant 4; this was less evident in the no cleaning treatment where the arena was not cleaned between exploration bouts (Figures 1, S1). Seeking to understand this unexplained tendency, we compared our trajectories with a study of *Lasius niger* by Khuong et al. [41]. They reported the following: “Regarding the statistics of exit heading direction, we observed a visible excess in favor of the lower right part of the canvas… We have no explanation for this bias so far, and it calls for further examination and testing.”

Their methodology does not specify whether there was arena cleaning between trials, and so it is possible that cuticular hydrocarbons influenced movement [17,47]. If this possibility can be excluded, Khuong et al. noted that some eusocial insects are sensitive to the magnetic field [48–51], and the movement bias in our data is in roughly the same south-westerly direction (Figure S1), though magnetoreception in *Temnothorax* seems improbable. A priori, one may have an expectation of a correlated random walk in a homogeneous environment, resulting in diffusive behaviour and a homogeneous exploration density at the macroscopic scale (e.g. [52]). What may be observed in Figures S1 and S2 is a switch between behaviours from intensive search in the central starting point (e.g. for the nest entrance in our data) to a stereotyped search (e.g. [53]). at which point the influence of a subtle environmental gradient, possibly interacting with intrinsic tendencies with respect to lateralisation, become evident in displacement direction. As with an MCMC method, given enough time, one would still expect all parts of the arena to be explored. Further review and testing across ant species should confirm whether heterogeneous exploration densities observed here are idiosyncratic or more widespread than previously recognised.

The experimental evidence for path avoidance of nestmates helped to inspire a new approach to tackling an analogous problem in the world of MCMC methods: slow convergence times owing to repeated sampling of the same region of probability space. The original MCMC method, Metropolis– Hastings, is well known for its random walk behaviour, yet is still a popular method because of its ease of implementation. We have shown that its efficiency can be substantially enhanced if it samples from the distribution 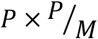, where *M* is a model (distribution) of where the MCMC walker has visited in the space, rather than simply *P* (the target distribution). The performance gain is particularly noticeable when sampling from sparse distributions, where the need to avoid revisiting previous parts of the space is especially acute. *T. albipennis* ants live in just such environments – where most of their surroundings hold little of relevance in terms of fitness, either food or potential nest sites – and hence such a mechanism should indeed be favoured by evolution. A shared substrate may serve effectively as a collective memory store, boosting the information-processing capability of the colony beyond the bottleneck of a single ant’s memory. Here we have just examined the enhancement to sampling performance for a single walker (ant) – of course in real colonies there will be many simultaneous walkers (ants), and their individual trajectories will be non-independent. More sophisticated MCMC methods that deploy independent parallel walkers, e.g. that presented by Wang and Landau [58], can be compared to our approach.

The model presented here retains a perpetual memory of where the ant has walked, although ongoing movement and pheromone deposition will lessen the relative weighting of past trails as pheromone accumulates elsewhere. In reality, pheromones will decay in strength over time. Although this is unavoidable, in a dynamic foraging environment it may well be preferable that repellent markers decay over time in any event, because they may quickly lose their relevance and even be maladaptive if they inhibit an ant from returning to an area where food has become available. Similar considerations may apply as with animals deciding how to weigh past observational memories with current ones; an exponential weighting is suitable in a Bayesian analysis [59]. It would be straightforward to introduce a reduction to the potency of the markers over time by multiplying the model by a decay factor, as in *e*^−*λt*^*M*, where *λ* is the decay constant. Theoretical consideration may be given to the optimal decay constant for different environments (or probability distributions), with sparse environments with a lower chance of opportunistic (randomly appearing) food favouring a lower constant, i.e. a slower decay. Such predictions may be readily tested in different ant species associated with different ecologies.

Theoretical consideration of self-organised behaviours such as ant collective foraging had predicted the value of inhibitory (negative) signals, before their empirical confirmation [60]. In certain circumstances they could make the foraging process much more efficient [61]. As anticipated, a ‘no entry’ signal has been found in Pharaoh’s ants *Monomorium pharaonis* to mark the unrewarding branch on a trail bifurcation. Concentrated at decision points – where the path branches – it could be used to complement attractive trail pheromones or to prevent an attractive pheromone becoming too strong through runaway positive feedback [62]. These ants are thought to have at least three types of foraging trail pheromone: one long-lasting attractive pheromone, thought to mark out the overall foraging network, and two short-lived attractive and repellent pheromones, to mark out short term food locations. The repellent pheromone was found to last more than twice as long as the short-lived attractive pheromone [63]. Simulations indicate that repellent pheromone increases the robustness and flexibility of the colony’s overall foraging response [64]. The usefulness of such negative or repellent markers has also been demonstrated in a swarm robotics context, in simulation [65,66], digitally on the robots themselves [67,68], or with real-world light-based markers [69–71]. Just as theoretical models of self-organisation predicted the value of negative markers before they were empirically confirmed, it is possible that advancements in Markov chain Chain Monte methods may inadvertently explain, or pre-empt the discovery of, certain behavioural mechanisms. Similarly, the biology of collective exploration is potentially a rich source of bio-inspiration for MCMC.

## Conclusion

Many organisms face the challenge of identifying and exploiting food sources in an unfamiliar and changing environment. A common pitfall in the search process is revisiting previously explored regions of space that do not contain anything of value. An evolutionarily ancient way of avoiding this problem is to leave an externalised ‘memory’ of the previous visit by depositing a chemical marker that can be used to make a quick decision not to re-enter that space when it is encountered later. For superorganisms like ants, that seek to maximise their foraging performance at the level of the colony, pheromone signals or cues such as cuticular hydrocarbon footprints may be used to coordinate the movement decisions of their nestmates (i.e. through stigmergy) such that an unfamiliar space is quickly explored. Here, we presented evidence for avoidance of the exploration of previous nestmates in *Temnothorax albipennis* ants, solitary foragers who would indeed particularly benefit from such a behaviour. We also demonstrated that a path avoidance behaviour enhances the exploration efficiency of the ants at the colony level.

In addition to empirical findings, after noting the analogy between ant exploration and sampling from unknown probability distributions, we developed an ant-inspired enhancement to Markov chain Monte Carlo methods, whereby a memory is kept of where a walker has moved through the probability space. Such a method was shown to significantly enhance the efficiency of the Metropolis–Hastings method [7,8] when sampling from sparse probability distributions, because the random walk type behaviour of the method is reduced, but at little computational cost in comparison to more advanced momentum-based methods such as Hamiltonian Monte Carlo [9]. When the ‘superorganism’ [43] is examined using our Bayesian framework [5,38], behavioural mechanisms such as individual trail markers become better understood in an ultimate sense [72] in relation to their facilitation of adaptive collective information-processing capabilities.

## Data accessibility

The *Temnothorax albipennis* ant data analysed here can be found at: http://dx.doi.org/10.5061/dryad.jk53j

The *Lasius niger* data analysed here is associated with [41] and can be found at: https://doi.org/10.1371/journal.pone.0076531.s005

## Author contributions

ERH drafted the paper, ran the simulations and analysed the data; NRF advised on social insect biology; RJB originated and advised on the theoretical concepts. All authors contributed to the present draft.

## Ethics statement

The ant *Temnothorax albipennis* is not subject to any licencing regime for use in experiments. The ants were humanely cared for throughout the experiment.

