## Supplementary Figures S1-6 for "The Bayesian Superorganism: externalised memories facilitate distributed sampling"

### Supplementary Information

**Figure S1.** We compared observations of heterogeneous arena exploration in our own ants with data on *Lasius niger* exploration. The number of ants leaving a 50x50cm region on each side is shown on the right. For our *T. albipennis* trajectories, the trajectory is coloured black until it first reaches the virtual 50x50cm boundary and blue thereafter (ants continue exploration rather than being removed as in the *L. niger* data). Not all ants in our data leave this virtual region, and so their final position after 45 minutes is marked instead. Statistics test a null hypothesis of uniform exit numbers on each side. Magnetic north is marked (although NB, local field can be affected by the building etc.)

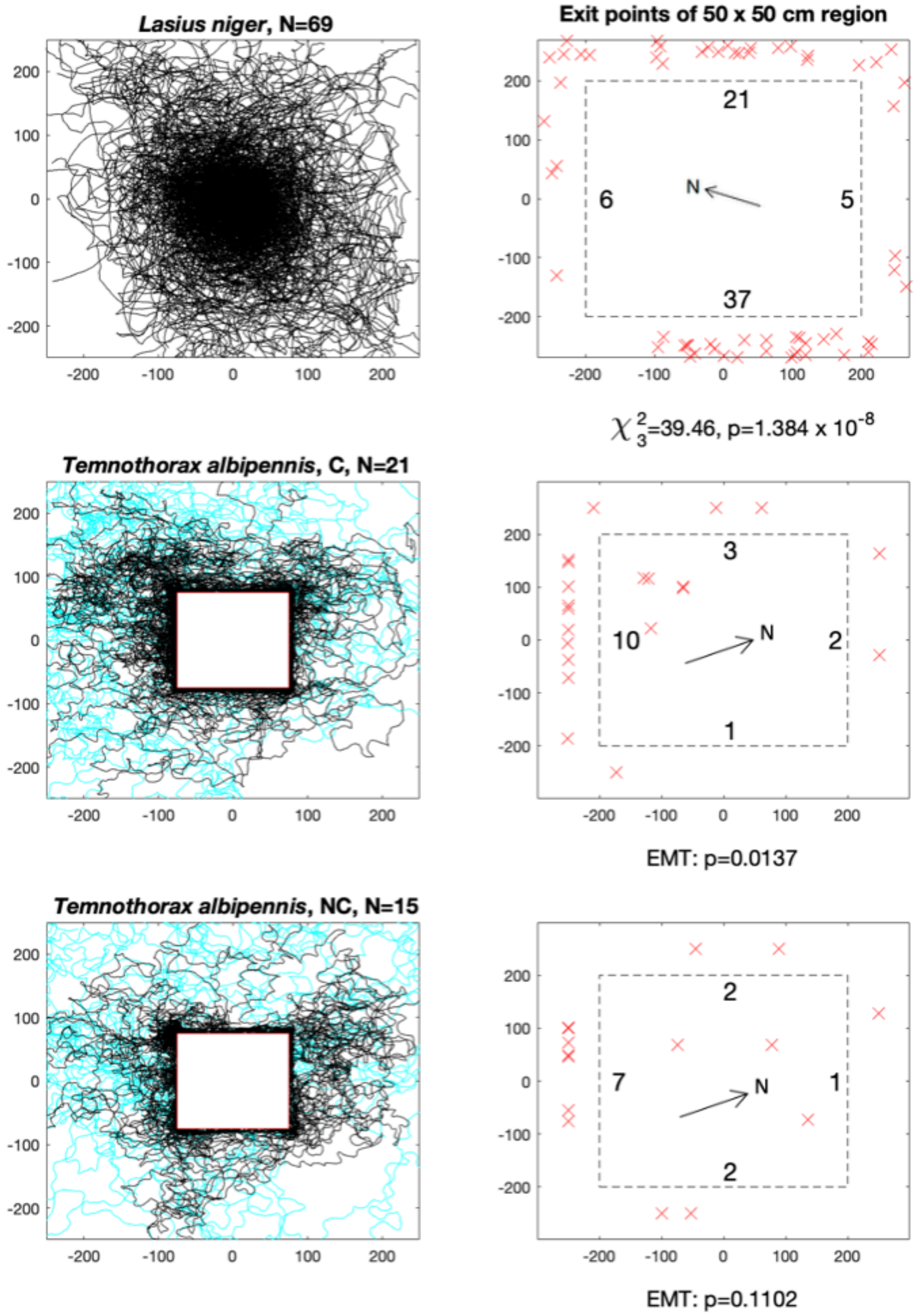

**Figure S2.** Distribution of ant positions during exploration (histogram bin counts), within a 12.5cm radius circle (i.e. diameter 25cm, half of 50cm virtual arena width), and outside that radius.

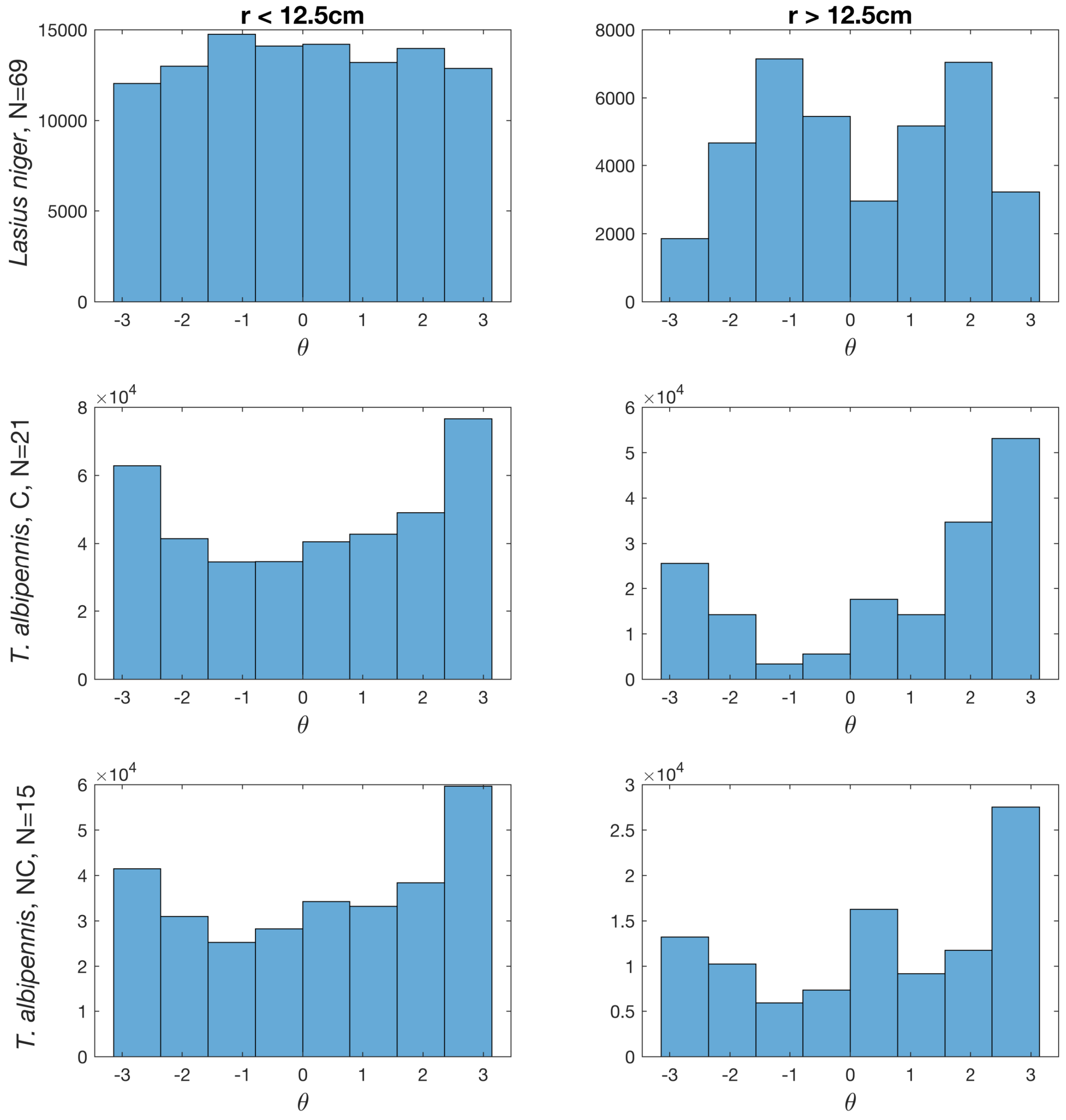


Although the *Lasius niger* trajectories from Khuong et al. uniformly explore the central region, their angular distribution is bimodal outside of it (c.f. Figure S1). The *Temnothorax albipennis* trajectories are somewhat uniform inside the radius (thigmotaxis on the square paper mask affects the distribution), and they are more obviously biased in the $-x$ direction ($\pi$ radians) outside this area. Note that the NC treatment angular distribution at $r > 12.5$cm is somewhat more uniform than the C treatment, especially around $[-\pi/2, 0]$.

Khuong, A., Lecheval, V., Fournier, R., Blanco, S., Weitz, S., Bezian, J. J., & Gautrais, J. (2013). How do ants make sense of gravity? A Boltzmann walker analysis of Lasius niger trajectories on various inclines. *PloS one*, *8*(10). https://doi.org/10.1371/journal.pone.0076531
**Dataset S1**, available at: https://doi.org/10.1371/journal.pone.0076531.s005

**Figure S3.** The individual trajectories of ants in the cleaning (C) treatment.

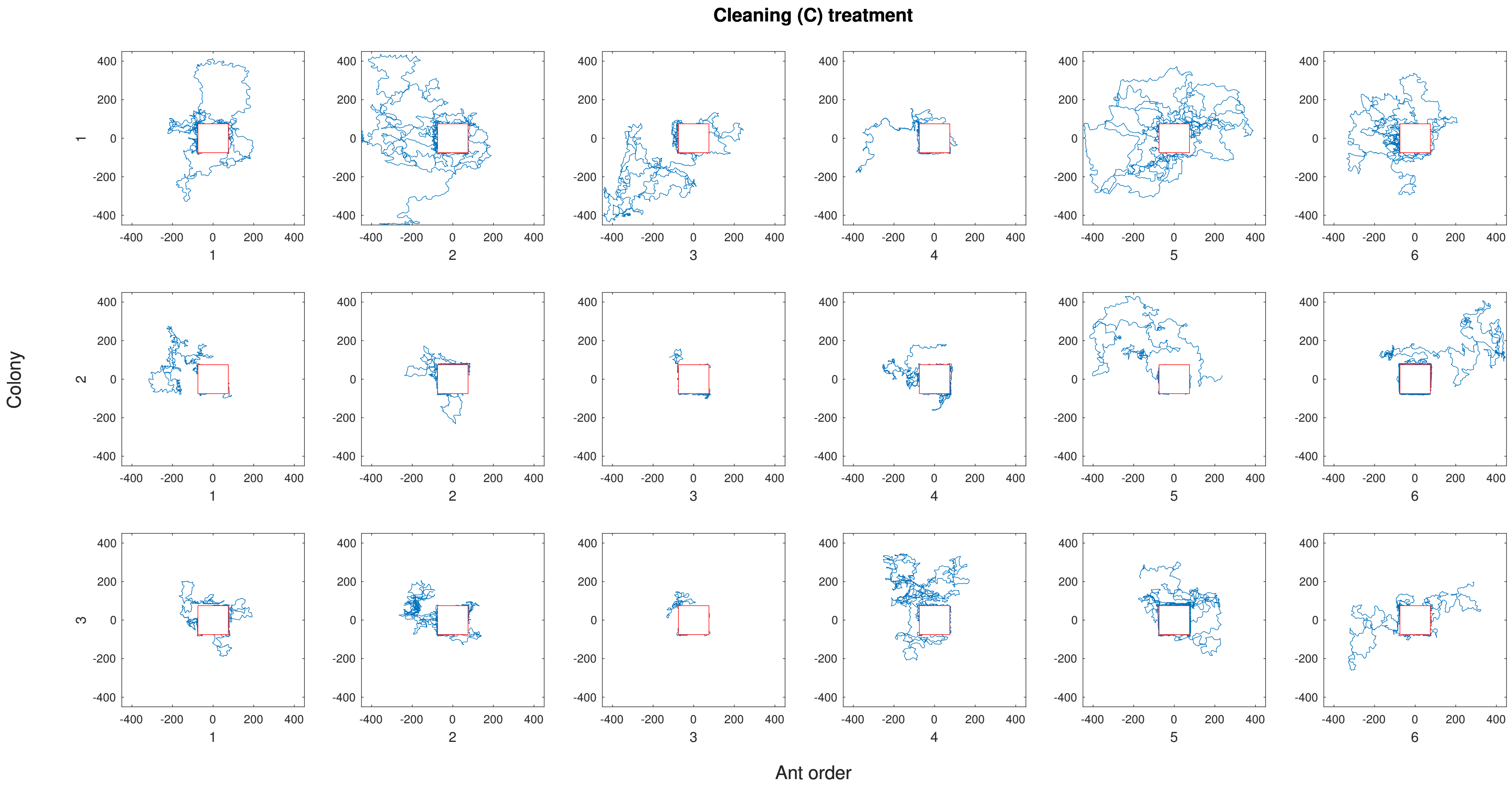

**Figure S4.** The individual trajectories of ants in the no cleaning (NC) treatment.

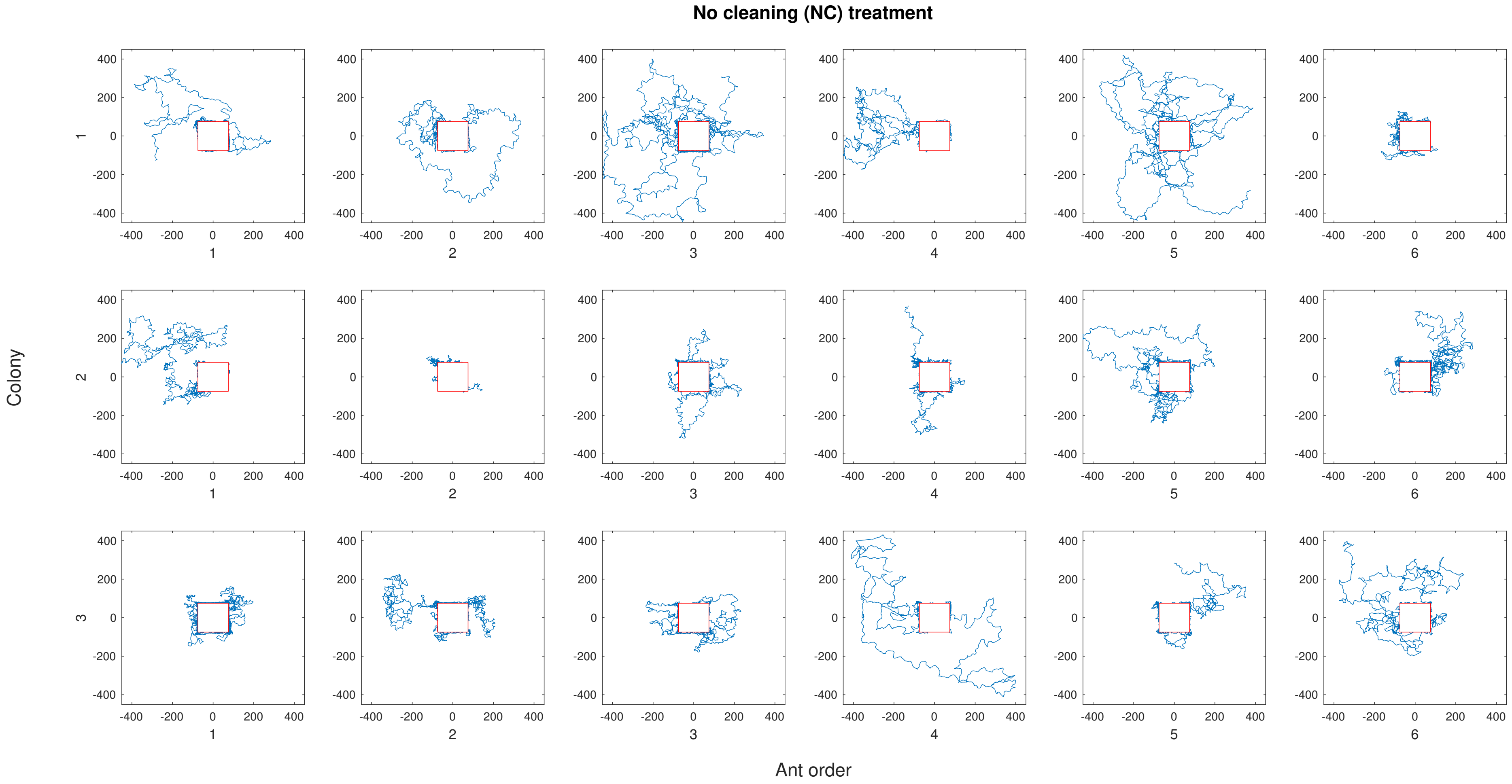

**Figure S5.** The cumulative trajectories of ants in the cleaning (C) treatment.

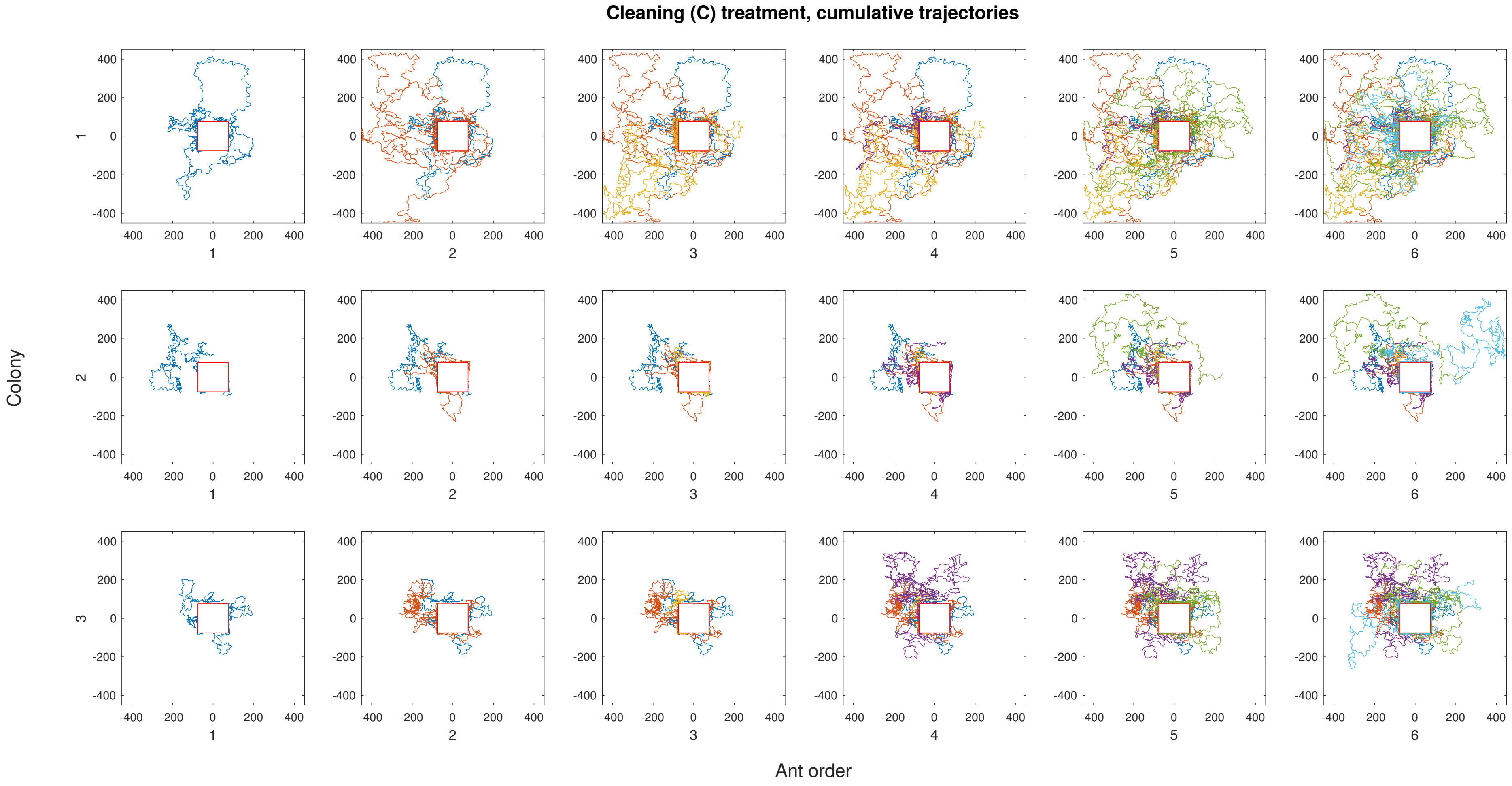

**Figure S6.** The cumulative trajectories of ants in the no cleaning (NC) treatment.

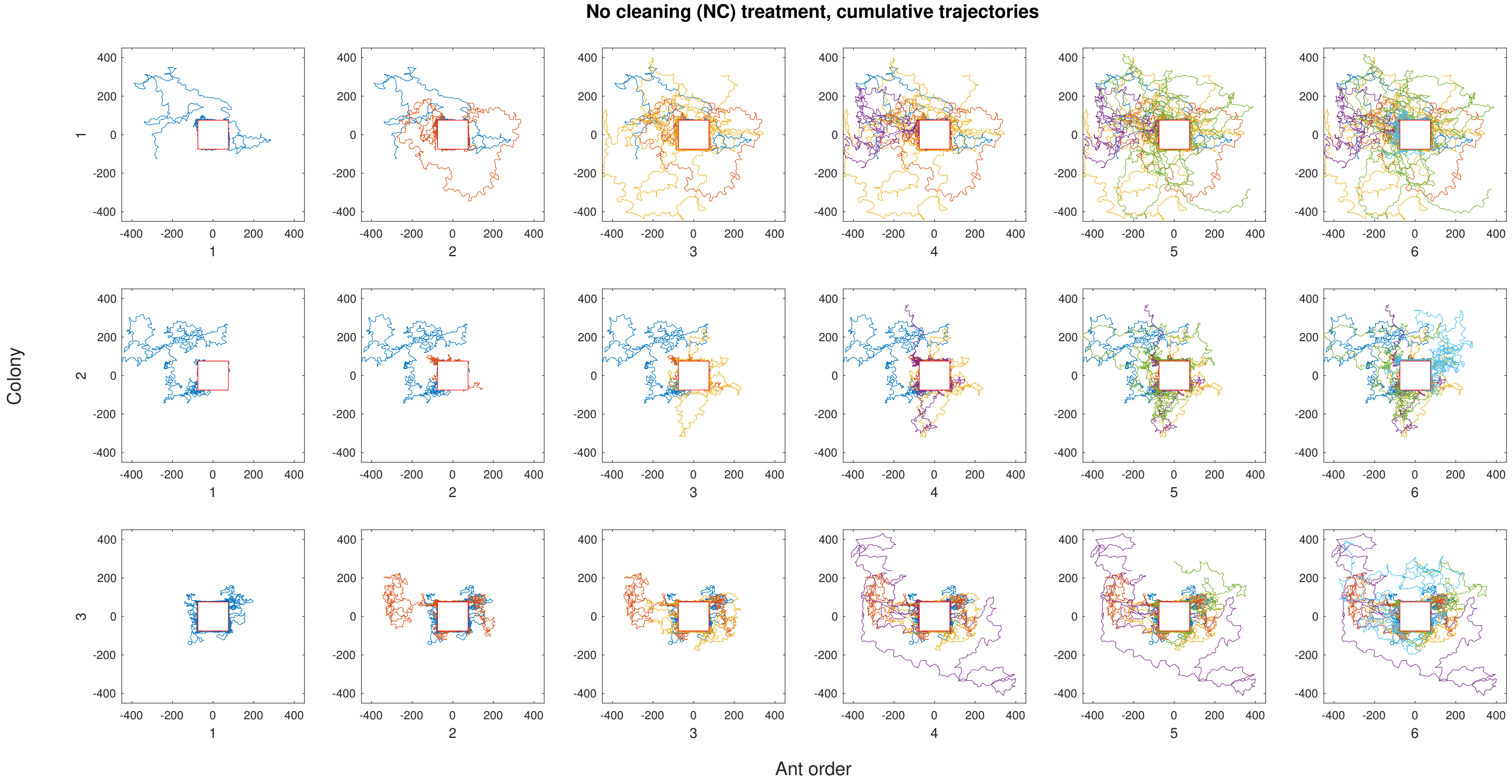
